# Quantifying the massive pleiotropy of microRNA: a human microRNA-disease causal association database generated with ChatGPT

**DOI:** 10.1101/2024.07.08.602488

**Authors:** K. Rowan Wang, Julian Hecker, Michael J. McGeachie

## Abstract

MicroRNAs (miRNAs) are recognized as key regulatory factors in numerous human diseases, with the same miRNA often involved in several diseases simultaneously or being identified as a biomarker for dozens of separate diseases. While of evident biological importance, miRNA pleiotropy remains poorly understood, and quantifying this could greatly aid in understanding the broader role miRNAs play in health and disease. To this end, we introduce miRAIDD (miRNA Artificial Intelligence Disease Database), a comprehensive database of human miRNA-disease causal associations constructed using large language models (LLM). Through this endeavor, we provide two entirely novel contributions: 1) we systematically quantify miRNA pleiotropy, a property of evident translational importance; and 2) describe biological and bioinformatic characteristics of miRNAs which lead to increased pleiotropy. Further, we provide our code, database, and experience using AI LLMs to the broader research community.

## Introduction

microRNAs (miRNAs) are small, non-coding RNAs that regulate major vital functions primarily by post-transcriptional suppression of gene expression. miRNA dysregulation has been linked to many complex diseases, including many cancers, Alzheimer’s disease, asthma, and cardiovascular disease ^1^. A single miRNA may regulate hundreds of genes and influence many distinct physiological processes ^2^. Consequently, dysregulation of a miRNA could result in multiple pathological perturbations to disparate biological pathways. For example, miR-21 has been shown to be a predictive biomarker for over 29 diseases ^3^. However, the number of diseases a given miRNA causally influences, what we refer to here as the miRNA’s *pleiotropy*, is an understudied characteristic of miRNAs in the literature.

Understanding miRNA pleiotropy has important implications for using miRNAs as biomarkers or as therapeutic targets. As argued by Jenike and Halushka, a miRNA with massive pleiotropy, like miR-21, lacks specificity and therefore makes a poor biomarker ^3^. A lack of specificity is also problematic for the burgeoning development of miRNA-based or miRNA-blocker therapeutics. Such a treatment may ameliorate one disease while causing other unintended iatrogenic conditions due to the miRNA’s broad influence on multiple different biological pathways.

At present, it is not known how many human miRNAs display pleiotropy on the same scale as miRNA-21. Being linked to 29 diseases, miRNA-21 may be an outlier among prevalent miRNAs, or it may be merely a representative case. Furthermore, if additional research is devoted to investigating miRNA-21, will it be found to be associated with an ever-increasing number of maladies? Or is there an expected upper limit? While of obvious importance to the field of translational miRNA biology, we ask and attempt to answer these questions for the first time herein.

Prior work has catalogued the effect of miRNAs on human disease through manual curation of causal miRNA-disease associations, notably in the Human-miRNA Disease Database (HMDD) ^4,5^. The HMDD has two potentially interesting versions: an earlier version (v3.2) which recorded only causal associations between a miRNA and a disease; and a larger, recent version (v4.0) which does not try to adjudicate causality, instead reporting causality-agnostic associations between a miRNA and disease. Both of these are quite large and represent a significant investment of expert person-hours to read and catalog miRNA manuscripts. With the increasing pace of miRNA research, and research publication in general, Artificial Intelligence (AI)-automated curation is an appealing option. Recent rapid advancements in the capabilities of Large Language Models (LLMs) have shown that these models are proficient at understanding and summarizing complex text, such as scientific literature ^6–8^. If LLMs could be employed with sufficient accuracy, the scalability of these systems makes them ideally suited to handle the increasing pace of scientific publication and aid in quantifying miRNA pleiotropy.

In this paper, we seek to quantify how many diseases a given miRNA causes and discover what intrinsic characteristics of a miRNA influence its pleiotropy. We leverage LLMs, specifically the Generative Pretrained Transformer (GPT) family built by OpenAI, to build a comprehensive database that maps out the causal landscape of miRNAs in human diseases. We describe the construction of this database, validate it against human annotators and the HMDD database (v3.2) for miRNA-causality, and apply it to further understand trends in miRNA research and miRNA pleiotropy. Through this endeavor, we hope to shed light on miRNA pleiotropy and provide a valuable tool for the broader research community.

## Methods & Data

To create the miRAIDD database, we first downloaded all miRNA-related abstracts from PubMed. We then annotated each abstract with whether the miRNA described therein was causally implicated in the disease tagged by the Medical Subject Headings (MeSH) terms of the abstract. From this, we created a database of miRNA-disease causal associations, where each causal link is supported by one or more research abstracts.

The overall methodology is illustrated in Figure 1.

**Figure 1:**
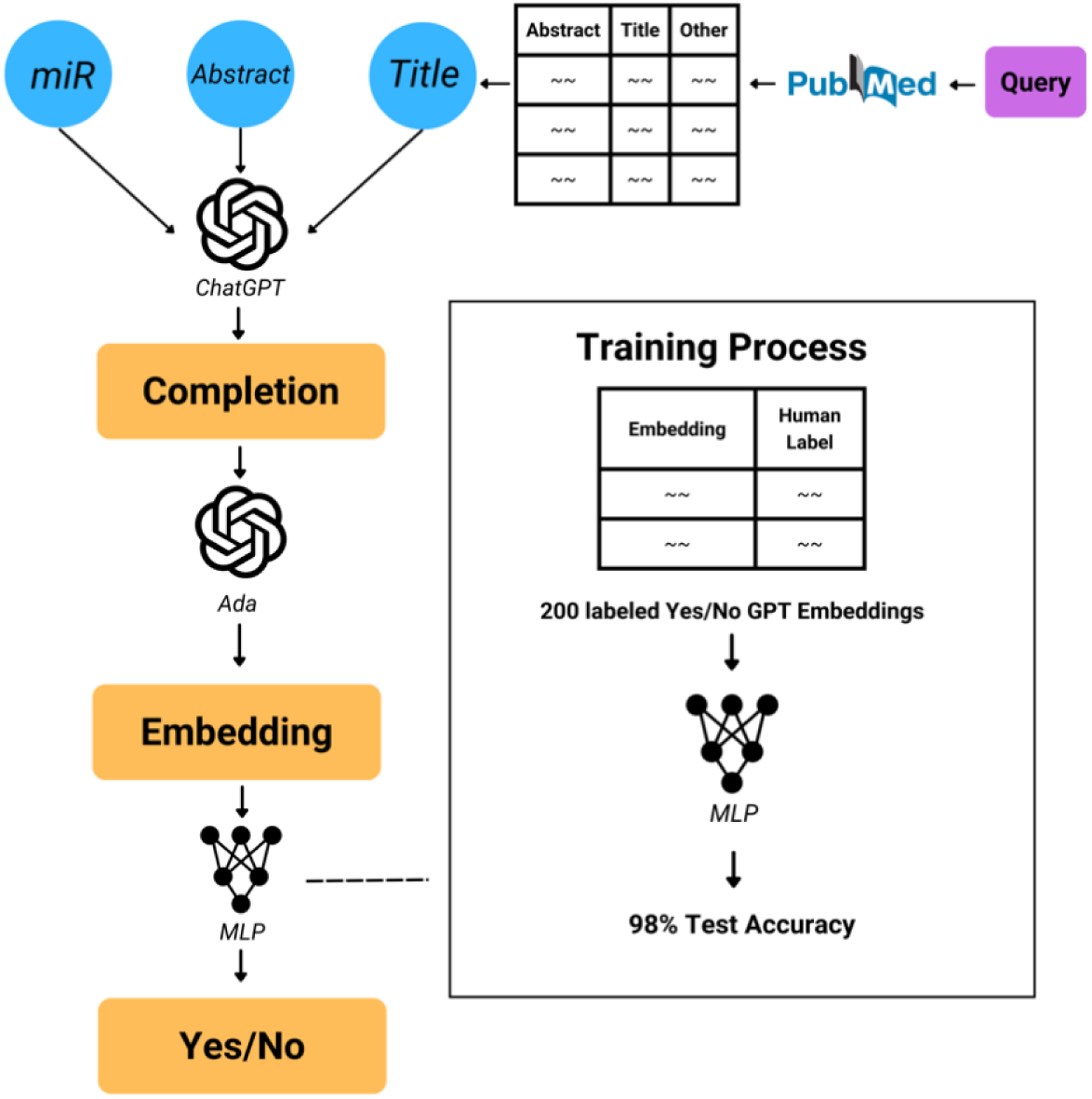
The miRAIDD pipeline.

### Pre-processing miRNA Abstracts

First, we compiled a list of all human miRNAs from miRBase (mirbase.com)^9^ We then searched PubMed for human-related abstracts for each of these human miRNAs. For this search we considered only the pre-miRNA, that is, the miRNA without the “-3p” or “-5p” appellation appended, which matched with both the pre-miRNA and mature miRNAs using PubMed.

The PubMed query was constructed as follows. We used the PubMed advanced search builder Application Programming Interface (API) together with PubMed’s Medical Subject Headings (MeSH) terms, with the query *“({miRNA-Name}[Title/Abstract]) AND (microRNA[MeSH Terms]) AND (Human[MeSH Terms]).”* This query was designed to retrieve papers that mentioned a specific miRNA in its title or abstract and contained the “microRNA” label and the “Human” label in their MeSH terms ^10^. We ran this query for all the pre-miRNAs in miRBase, instantiating *{miRNA-Name}* to a unique miRNA each time. This resulted in a list of 85,454 miRNA-abstract pairs. This set of miR-abstract pairs included multiple instances of the same abstract, in the cases where more than one miRNA was described in a single abstract.

Finally, to identify the disease(s) associated with each abstract, we considered the paper’s MeSH terms together with the Human Disease Ontology (DO) ^11^. First, we retrieved the abstract’s MeSH terms tagged with “Pathology” using the MeSH SPARQL API ^12^. To limit this to a codified ontology of human diseases, we used the intersection between the “pathology” MeSH terms and the disease terms in the DO in the following manner. For every MeSH term, if that term was exactly in the DO, then we used that term as the “disease” of the abstract. If not, we looked upward from the MeSH pathology term in the MeSH terminology tree at the ancestors of that term, and used the first ancestor to match with a disease in the DO as the disease for the abstract. This process was repeated for each pathology MeSH term in the abstract. If no matches were found in the DO for an abstract, then the abstract was removed from further consideration.

### Assessing miRNA-Disease Causality

Next, we used ChatGPT to analyze whether each abstract provided evidence that its given miRNA played a causal role in a disease. To do so, we used a one-shot prompt, where we first listed general characteristics of abstracts that might provide causal evidence and then included an example abstract and completion from Nan et al. ^13^

The prompt was finally expanded with the text of the Title, Abstract, and the name of the miRNA for each query to ChatGPT. To be clear, each prompt includes the full text of the example abstract by Nan et al., which did not change, together with the full text of the actual abstract to be analyzed. The full prompt format Is available in the Supplementary Appendix.

Output from ChatGPT were, on average, 40 (±15) words long. A representative example follows, which ChatGPT produced in response to the abstract by Zhang et al.^14^:

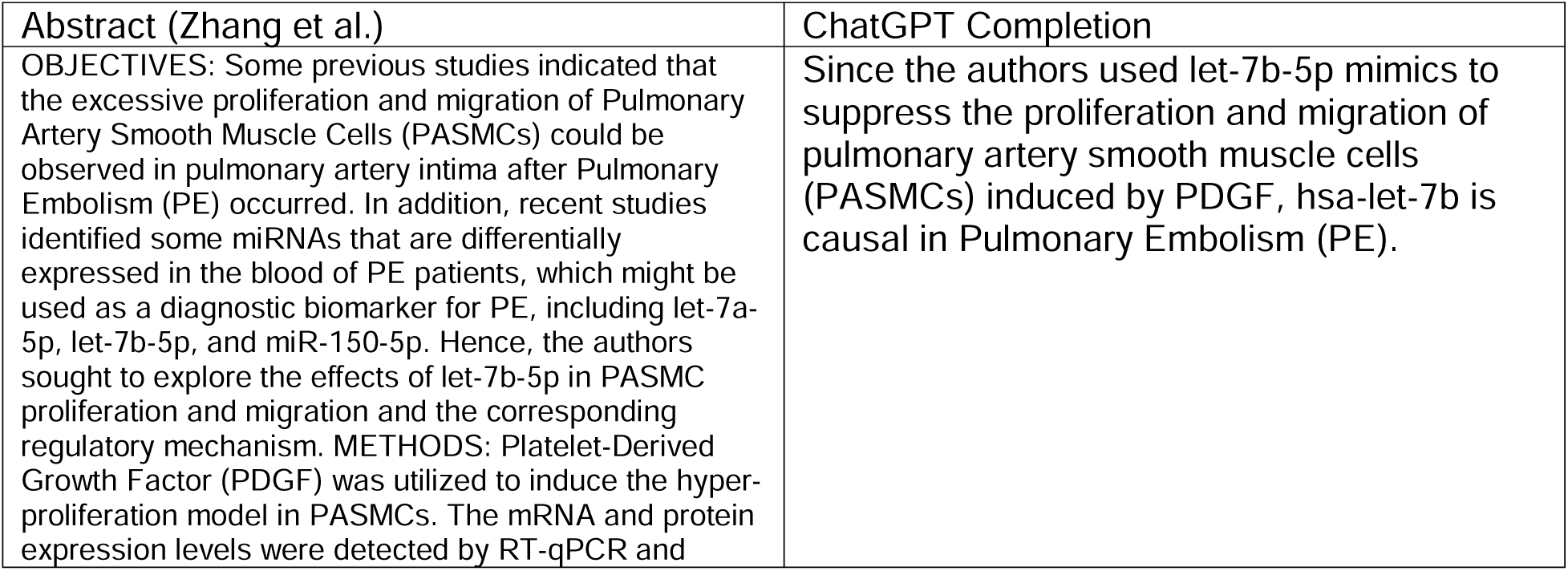
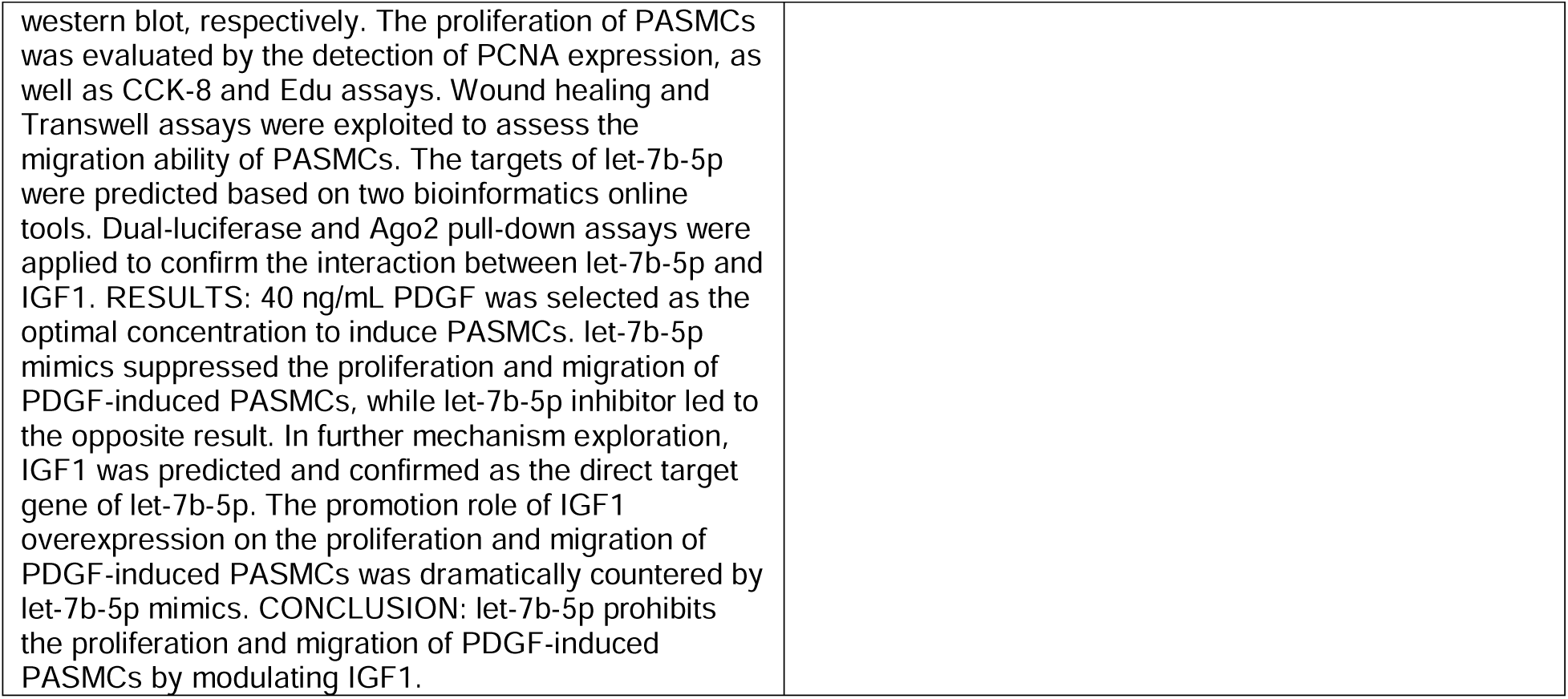

These responses are reasonable, but a human annotator would still be required to read each of them to determine if ChatGPT was saying that yes, there was causality, or no, there was not. Reading all 85,454 AI-generated responses would still represent a substantial investment of human effort, one that we wished to avoid.

Accordingly, we used a machine-learning algorithm to convert ChatGPT’s textual responses into binary yes/no answers that indicate miRNA-disease causality. To do so, we embedded ChatGPT’s responses with OpenAI’s ada model (text-embedding-ada-002) using default parameters, yielding 1536-dimensional embeddings ^15^. Two of the present authors manually annotated 250 randomly-selected textual responses as either stating that “yes” there was a causal link, or “no” there was no evidence of causality between the miRNA and disease in the abstract. After annotation, these authors discussed their annotations and came to agreement on two instances of disagreement. Using 200 of this labeled training data, we then built a two-layer Multi-Layer Perceptron (MLP) to identify whether the embedded textual responses were causal, leaving 50 for testing. This MLP had a hidden dimension of 128 and was trained over 500 epochs with a learning rate of 0.01 using PyTorch ^16^. We then ran this on the 85,454 textual responses from ChatGPT embedded with the ada model.

### Composition of miRAIDD

We composed the miRNA AI Disease Database (miRAIDD) from the PubMed abstracts, the MeSH terms, ChatGPT output, and causal label. More specifically, for each abstract-miRNA pair, we include: the text of the abstract; the title; the date published; the list of MeSH terms annotated to the abstract in PubMed; the PubMed ID; the miRNA name; the full ChatGPT completion in response to the prompt; the causality label from the MLP; and the diseases from DO. The miRAIDD is freely available for download at https://github.com/Wanff/miraidd.

### Additional miRNA Data Sources: the Intrinsic miRNA and Pleiotropy Properties (IMAPP) database

We also combined the number of diseases each miRNA caused with other public data sources into a single database for use in our analyses. These factors are shown in Table 1 and included the maximum and average expression levels, number of tissues present (miRMine ^17^), validated targets, predicted targets from three different miRNA-target providers (miRanda, TargetScan, MirDB; accessed through multiMiR ^18–21)^, evolutionary conservation, year discovered (miRBase ^9^), and percentage cancer (miRAIDD).

**Table 1:** IMAPP

| Name | Description | Source |
| --- | --- | --- |
| miR name | Name of the miRNA | miRBase |
| Diseases | List of diseases the miRNA causes | miRAIDD |
| Pleiotropy | Number of diseases the miRNA causes | miRAIDD |
| Number of Papers | Number of papers published about the miRNA | miRAIDD |
| Conservation | Sum of miRNA family size | miRBase |
| Average Expression | Average expression of the miRNA in public datasets | miRmine |
| Max Expression | Maximum expression of the miRNA in public datasets | miRmine |
| Number of Tissues | Number of different tissues where miRNA has been expressed | miRmine |
| Year Discovered | Year of seminal miRNA publication in miRBase | miRBase |
| Percent Cancer | Percent of diseases the miRNA causes that are cancer-related | miRAIDD |
| Validated Targets | Number of Validated Targets | multiMiR |
| miRanda Targets | Gene targets predicted by miRanda algorithm | multiMiR |
| TargetScan Targets | Gene targets predicted by TargetScan algorithm | multiMiR |
| miRDB Targets | Gene targets predicted by miRDB algorithm | multiMiR |

To obtain lists of validated and predicted targets, we used mulitMiR to retrieve results from miRanda, TargetScan and MiRDB, which were considered as three separate variables. Due to formatting differences, we first queried multiMiR with a pre-miRNA (without –3p and –5p) name. If this returned no results for a particular target database, we then queried multiMiR with both the –3p and –5p mature miRNA names, summing the results together. Queries were conducted including both conserved and non-conserved target sites in target genes. The top 20% of genes returned were retained as high-confidence targets. MultiMiR was also used to obtain a count of validated gene targets, which are published, experimentally validated miRNA-target interactions.

To obtain expression levels and number of tissues expressed, we queried miRMine using the pre-miRNA name to get a list of expression values and tissues associated with each pre-miRNA. If no expression values were found, we imputed a value of 0.

We computed miRNA evolutionary conservation as the sum of the miRNA family size in miRbase, following our previous work ^22^. This is a count of the number of species where the miRNA has been observed without sequence changes, and was previously found to be predictive of disease causality.

The year a miRNA was discovered is set to be the date of the seminal publication in miRBase about that miRNA. Percentage cancer is calculated to be the percentage of diseases caused by the miRNA that contain cancer-related words in their name (lymphoma, carcinoma, etc.) in miRAIDD. The full list of qualifying cancer-related terms can be found in the Supplementary Appendix.

The above data were combined with the miRNA’s list of diseases caused, pleiotropy count, and number of papers mentioning the miRNA from the miRAIDD into our Intrinsic miRNA and Pleiotropy Properties (IMAPP) database, which is freely available for download at https://github.com/Wanff/miraidd.

### Statistical Tests and Analysis

Statistical analysis related to the pleiotropy of miRNAs was performed in R and Python. Regression analysis was conducted using the lm() function in R Version 4.2 ^23^.

Significant covariates were declared based on a coefficient not equal zero with probability p < 0.05. Uniform Manifold Approximation and Projection (UMAP) clustering was carried out with the Python umap-learn package using standard hyperparameters, with number of neighbors set to 15 and minimum distance set to 0.1 ^24^. Correlation and partial correlation matrices were generated with the pingouin Python package (Version 0.5.3) with standard hyperparameters ^25^.

## Results

### MiRAIDD

To validate the miRAIDD pipeline, we validated each of the pipeline’s components individually. First, we found that the final MLP at the end of the pipeline, which converts the embeddings of ChatGPT’s responses to binary labels, achieved 98% accuracy on a test set of 50 labels.

Next, we validated our whole AI-based causality determinations against human researchers and HMDD. We randomly chose 100 abstracts from our PubMed search and had three human researchers with miRNA experience read and determine if evidence of a miRNA causing a disease was given in the abstract. On average, the human judges agreed with each other 78.6% of the time, while our ChatGPT-based pipeline agreed with the human judges 76.3% of the time (Table 2).

**Table 2:** ChatGPT vs Human Pairwise Agreements

|  | Human 1 | Human 2 | Human 3 | ChatGPT |
| --- | --- | --- | --- | --- |
| Human 1 | - | 81% | 78% | 80% |
| Human 2 | 81% | - | 77% | 71% |
| Human 3 | 78% | 77% | - | 78% |
| ChatGPT | 80% | 71% | 78% | - |

We also compared our miRAIDD causality determinations to HMDD v3.2: a manually curated miRNA-disease causal association database ^5^. Of the 19,017 miR-abstract pairs in the intersection between the two datasets, we found that miRAIDD agreed with HMDD’s labels 77.9% of the time ^4^, representing broadly similar agreement to our own human judges. Despite this similar performance, HMDD and miRAIDD used slightly different criteria to determine disease-causality. HMDD excludes papers in which miRNA interventions enhance drug effects but don’t modulate disease state directly, while miRAIDD, according to our AI prompt, doesn’t necessarily do so ^26^. MiRAIDD also did not consider review articles, while these were considered evidence in some instances in HMDD. Supplemental Table S1 shows some of the specific disagreements between miRAIDD and HMDD.

MiRAIDD was generally larger than HMDD v3.2, where HMDD has 35,547 annotations (miRNA-abstract pairs), 19,281 unique papers, and 5,630 unique papers that demonstrate a causal miRNA-disease association. In contrast, miRAIDD has 85,454 annotations, 51,410 unique abstracts, and 23,346 unique abstracts that demonstrate a causal miRNA-disease association.

### Intrinsic Properties of miRNA

Our primary goal in creating the miRAIDD was to quantify the pleiotropy of miRNAs and identify the intrinsic characteristics of a miRNA which affect that pleiotropy. To do so, we combined miRAIDD, which provided measures of pleiotropy and research effort, with other publicly available miRNA databases to identify factors influencing the pleiotropy of miRNAs, as described above (Methods, Table 1), into the Intrinsic miRNA and Pleiotropy Properties (IMAPP) database. IMAPP is available along with miRAIDD at https://github.com/Wanff/miraidd.

First, we visualized the IMAPP data using the UMAP technique. We found that miRNAs seemed to separate into two main groups (Figure 2), one corresponding to cancer-causing miRNAs and the other to non-cancer-causing miRNAs ^24^. This is indicative of the predominance of cancer-based research on miRNAs, a trend previously noted, ^27^ and which has previously required bioinformatic corrections to address ^28^. We noted the outlier at bottom left (Figure 2) represents five miRNAs that cause no diseases and have few non-zero values in IMAPP. These were retained in further analysis to avoid undue bias against understudied miRNAs.

**Figure 2:**
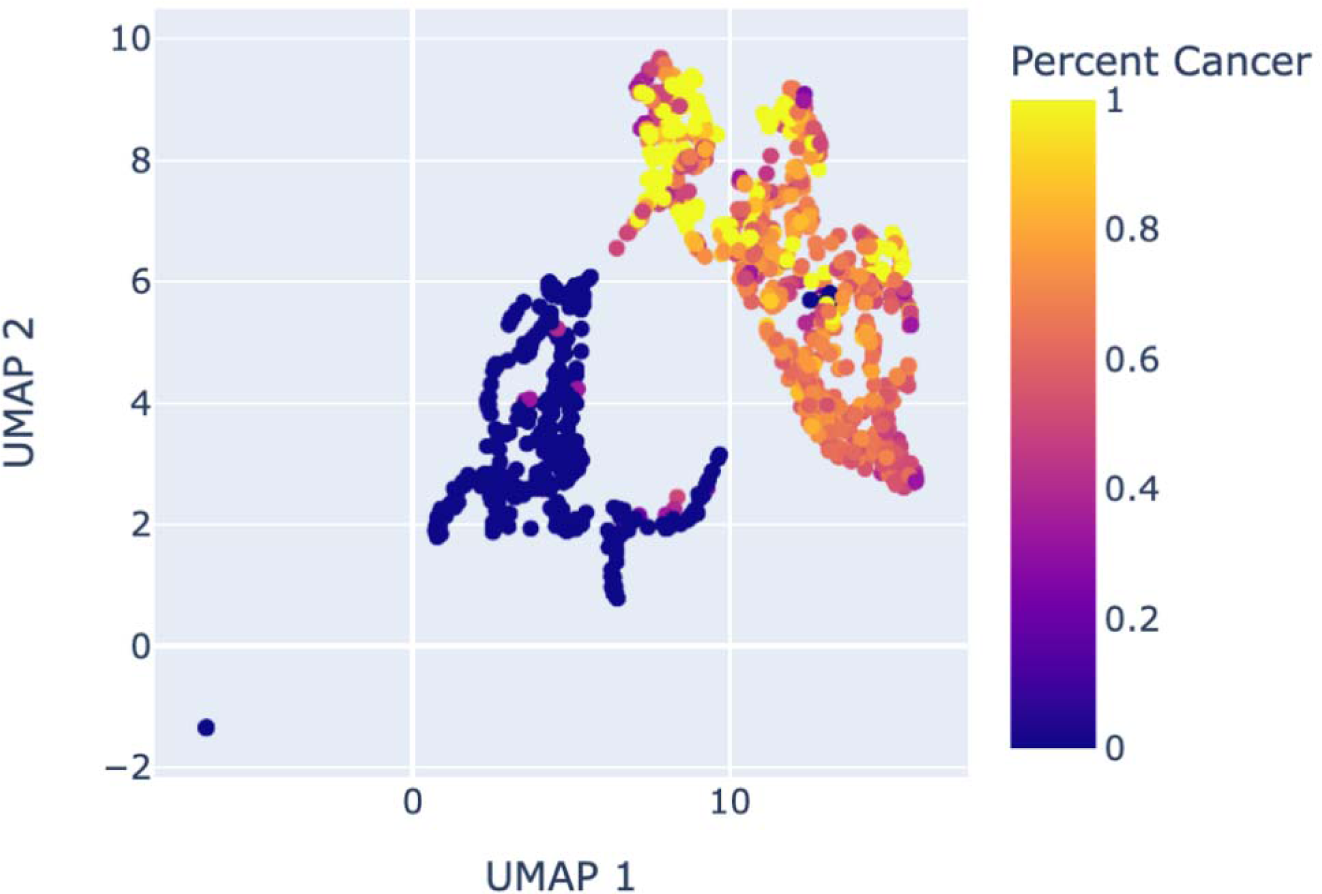
Two-dimensional UMAP on the IMAPP miRNA data

### MiRNA Pleiotropy

Next, we note that research effort drives miRNA pleiotropy (Figure 3). Intuitively, each miRNA-disease causal association is linked to a set of published papers, so the more diseases a miRNA has been shown to cause, necessarily, the more papers that have been written about it.

**Figure 3:**
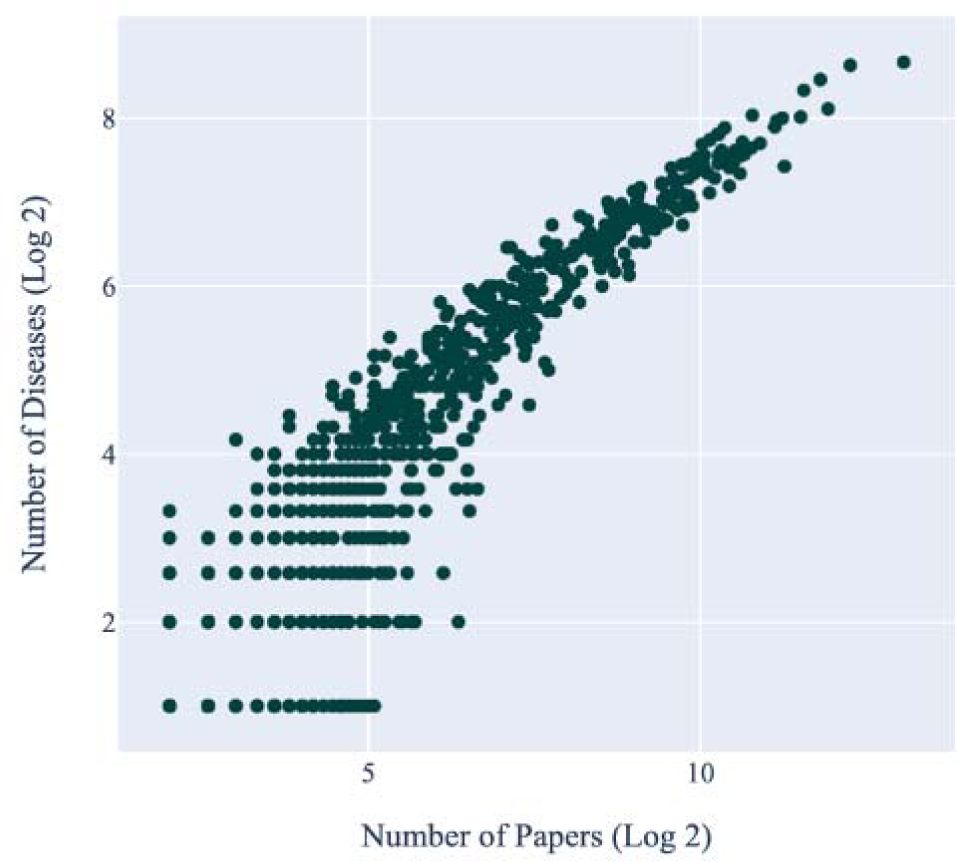
Number of papers written about a given miRNA and the number of diseases it has been shown to cause. Log-log scale.

In fact, the number of papers written about a miRNA is the single biggest predictor of miRNA pleiotropy. Predicting miRNA pleiotropy from research effort alone using standard linear regression yields an r^2^ of 0.733, whereas predicting pleiotropy from all the miRAIDD and IMAPP factors collectively was only somewhat better (r^2^ = 0.876). To some extent, this is a consequence of how we have measured the pleiotropy of miRNAs: the number of abstracts presenting evidence of unique disease causality is the number of diseases the miRNA causes. Below, we show a correlation matrix that illustrates the strong correlation of research effort with pleiotropy (Figure 4, right). Interestingly, in this correlation matrix, we see that several intrinsic factors of miRNAs are strongly correlated. For example, both miRNA average and maximum expression are strongly correlated. We also see that two of the miRNA gene target metrics, miRDB and miRanda, are correlated with each otherTargetScan.

**Figure 4:**
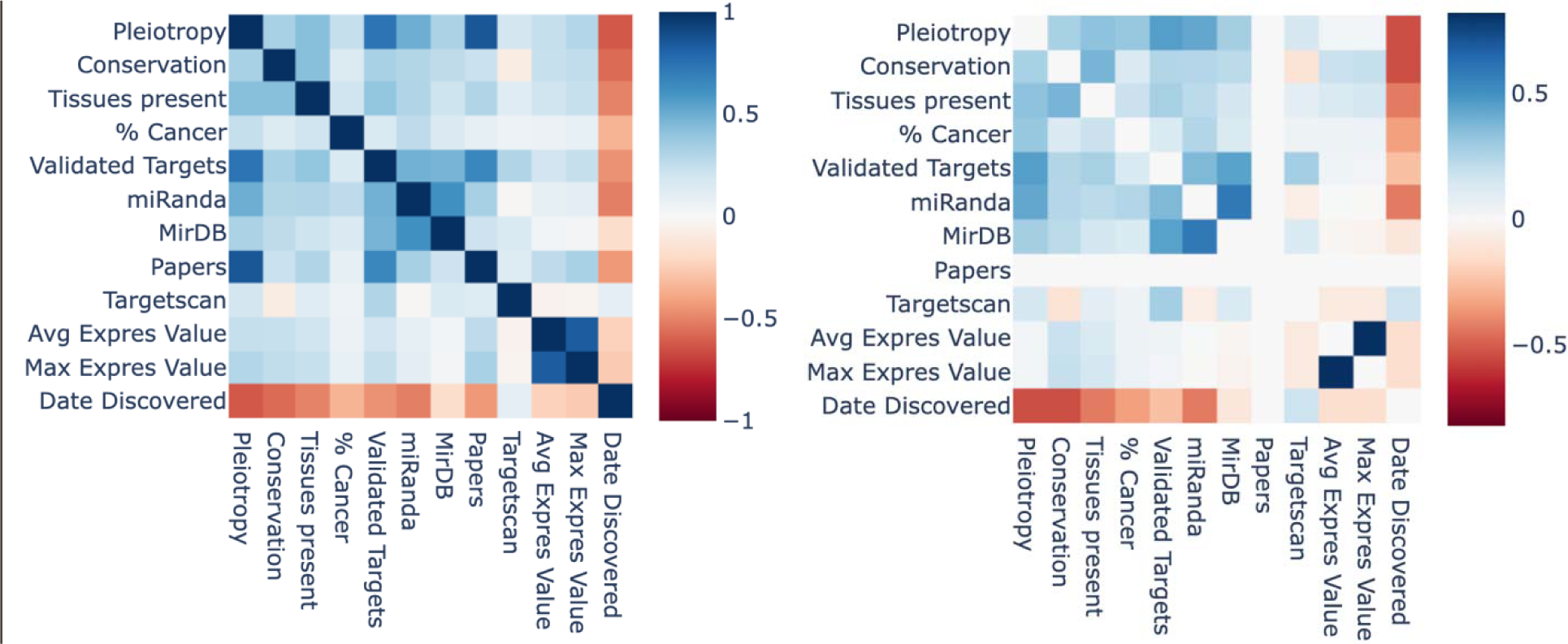
Correlation matrices showing standard Pearson correlation between different biological factors, with clustering. Left: Normal correlation matrix Right: Adjusted for number of papers.

However, to answer our original question about which intrinsic characteristics of a miRNA affect its pleiotropy, we adjust for research effort. We show a partial correlation matrix adjusted for papers in Figure 4 (left). When compared to the unadjusted correlation matrix (Figure 4, right), the correlations between each feature and pleiotropy are in the same direction but generally lower in magnitude. We then performed a regression analysis of intrinsic miRNA features on pleiotropy while adjusting for number of papers, showing some significant predictive performance (r^2^ of 0.370). Since average and maximum expression were tightly colinear, we removed average expression from this regression. The most predictive features were the number of validated targets and miRandaTargetScan targets, percent cancer, number of tissues where the miRNA is present, and conservation (Table 2). MiRdb targets were also predictive, although correlated with miRanda targets, and less predictive than miRanda. Interestingly, TargetScan targets were both predictive of and anti-correlated with pleiotropy after adjusting for number of papers.

**Table 2:** Linear Regression on Pleiotropy, Adjusted for Research Effort

| Feature | P-Value | Effect Size (normalized) |
| --- | --- | --- |
| Conservation | 6.66e-4 | 1.18 |
| Max Expression | 0.330 | -0.21 |
| Number of Tissues Present | 3.14e-8 | 2.09 |
| Number of Validated Targets | <2e-16 | 4.72 |
| Number of miRanda Targets | <2e-16 | 4.25 |
| Number of miRdb Targets | 0.0023 | 1.12 |
| Number of TargetScan Targets | 0.0014 | -1.23 |
| Year Discovered | 0.646 | 0.02 |
| Percent Cancer | 8.78e-15 | 2.27 |

Many miRNAs exhibit substantial pleiotropy, rendering them of little use as biomarkers. Specifically, we found 64 miRNAs cause 64 or more diseases. A table of these massively pleiotropic miRNAs can be found in Supplemental Table S2.

Given that there are many miRNAs with exhibiting strong pleiotropy, we may well ask: is pleiotropy bounded? Will miRNAs such as miR-21 eventually be shown to cause almost all diseases? We sought to answer this question by analyzing trends in miRNA research over time, considering the miRNAs with the highest pleiotropy and the most papers written, such as miR-21 (Figure 5). We looked at trends among miRNAs that are in the 95^th^ percentile for both number of diseases and number of papers (n = 57). For 93% (53 out of 57) of these miRNAs, which we term here the “popular” miRNAs, the number of new miR-disease causal associations per year peaked on or before 2013, indicating that the rate of discovery of new diseases caused by each of these miRNAs is decreasing (Figure 5, green bars, orange line).

**Figure 5:**
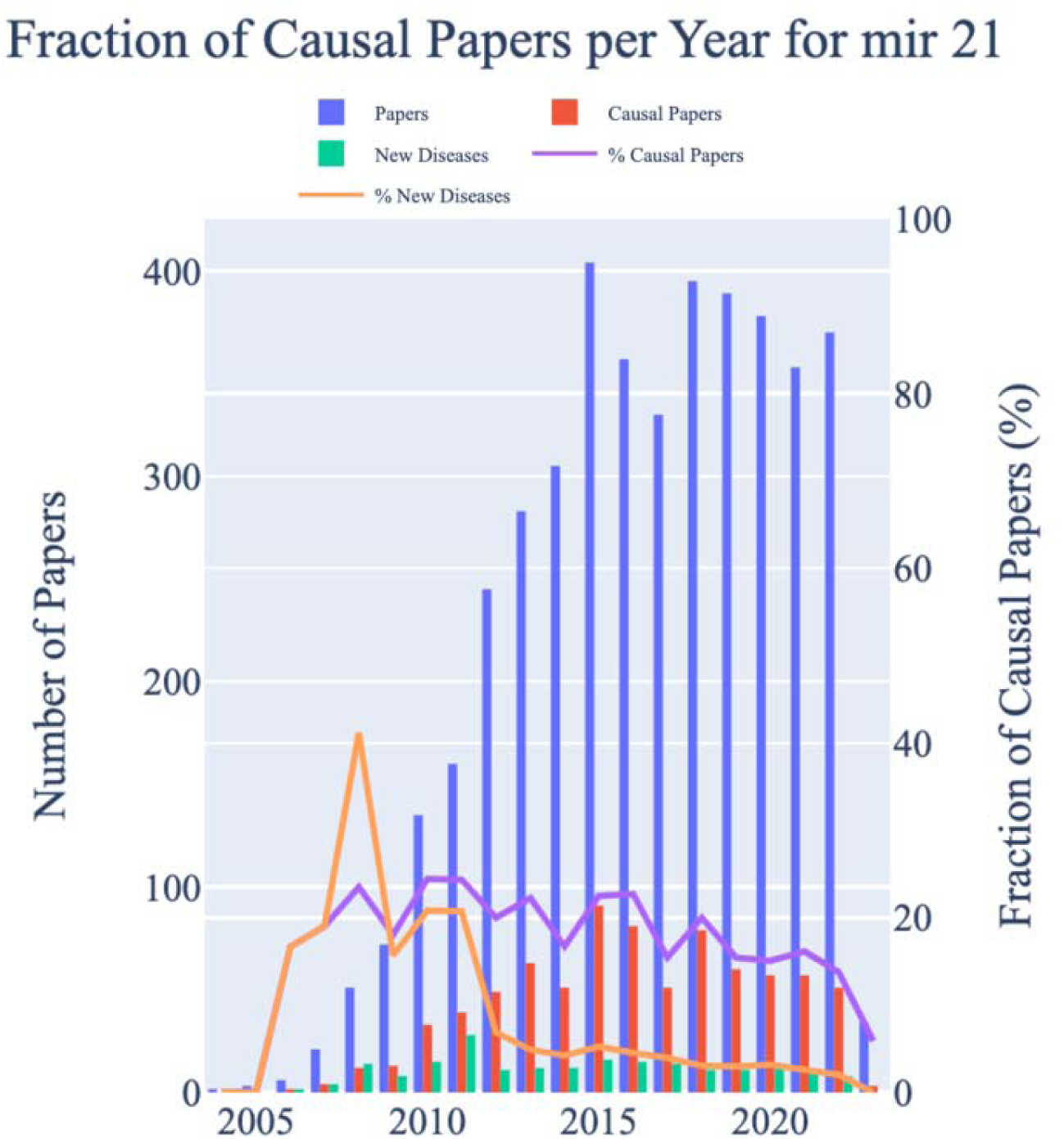
Trends in research on miR-21 over time. % Causal Papers represents the percentage of papers written about miR-21 that year that presented causal evidence for a miR-disease link (purple). % New Diseases represents the percentage of papers that discovered a new miR-21-disease causal association (orange).

We attempted to estimate the rate of decline in new diseases for each popular miRNA by fitting a line to these data. We chose to analyze data from 2012 to 2020, choosing 2012 as a date by which miRNA research seems to have been firmly established, and limiting ourselves to data prior to 2021, before research slowdowns caused by the COVID-19 pandemic. Then, for each popular miRNA, we fit a line for the fraction of papers that discover a new miR-disease causal association from 2012 to 2020. For the 24 miRNAs that had statistically significant trends (p-value < 0.05), all of them were negative, indicating fewer new diseases being attributed to the miRNA over time. While this does not show that new diseases caused by massively pleiotropic miRNAs such as miR-21 will cease to be discovered, it does show that such discoveries are becoming less frequent.

miRAIDD also enables us to discover miRNAs that are potentially understudied in the literature. To do so, we searched for miRNAs that our model predicts should cause more diseases than they are currently known to, while having relatively few papers (Supplemental Table S3). Many of these miRNAs are homologs or family members of other miRNAs with established pleiotropy, such as miR-451-b or hsa-let-7c. Others have very few papers, making it difficult to confidently estimate the difference between observed and expected pleiotropy. Some notable understudied miRNAs, still with many publications, are shown in Table 3.

**Table 3:** Notable Understudied miRNAs

| miRNA | Predicted Pleiotropy | Actual Pleiotropy | Number of Papers |
| --- | --- | --- | --- |
| hsa-mir-369 | 21 | 14 | 57 |
| hsa-mir-1915 | 13.5 | 6 | 32 |
| hsa-mir-522 | 24 | 17 | 36 |
| hsa-mir-1301 | 33 | 26 | 63 |

## Discussion

In this paper, we have developed a comprehensive database of human miRNA-disease causal associations with ChatGPT. Using this database, we analyzed trends in miRNA research and pleiotropy. We quantified miRNA pleiotropy systematically, confirming Jenike and Halushka’s observation that miR-21, and other miRNAs, exhibit massive pleiotropy in the literature. We found miR-21 caused diseases ranging from diabetes to leukemia to asthma.

While miRNAs with substantial pleiotropy may make for dubious biomarkers, their therapeutic potential is brighter. Theoretically, a pleiotropic miRNA could ameliorate certain disease states when upregulated but also aggravate others. However, it is also possible that a balanced expression level of a pleiotropic miRNA could lessen adverse effects systemically and restore homeostasis. Indeed, translational work with pleiotropic miRNAs proceeds with the clinical trial NCT02581098, studying the effects of miRNA-21 therapy on diabetes.

Among the factors predictive of miRNA pleiotropy were the number of validated and predicted targets by multiple target providers. After adjusting for the number of published papers, the targets predicted by the Miranda algorithm emerged as the most predictive. Interestingly, miRDB, a machine learning-based target prediction algorithm, was not nearly as predictive after controlling for number of papers. This effect might be explained by miRDB’s training process, which utilized existing literature, while Miranda and TargetScan, both of which are based primarily on the miRNA sequence and other genetic information, provide literature-independent insights. Interestingly, TargetScan was negatively correlated with miRNA pleiotropy, and also negatively correlated with miRanda, miRNA expression, and conservation. This may indicate that TargetScan is doing something decidedly different from other research in miRNAs, which may be a good thing for new hypothesis generation, or a bad thing for validation of accepted knowledge.

The implications of automated knowledge generation using powerful AI systems are immense. We used ChatGPT to scalably and efficiently generate knowledge from preexisting literature, and in the future, similar databases could be generated quickly in a variety of domains. Our particular application is an attractive one for AI automation, since 1) the task is prohibitively time-consuming to perform with expert humans; 2) AI models are already roughly as accurate as humans; and 3) the presence of errors is expected and leads only to slightly noisier data, rather than more serious consequences in medical applications ^29^. Indeed, others have shown successes using LLMs and generative AI for identifying biological relationships from microbiome literature ^30^.

However, limitations remain. ChatGPT’s classification accuracy of miRNA-disease causation from an abstract is imperfect, although similar to human classification accuracy; in the future more advanced AI models could be used, or ones with more specific domain training. Other sources of error in our pleiotropy estimates arise from potential redundancy in the disease ontology, potential omissions when indexing PubMed for relevant papers, and inconsistent nomenclature for miRNAs that has changed over time and across experiments and packages.

Additionally, automated extraction of concrete scientific knowledge from published abstracts inherent challenges, such as publication biases towards positive results ^31^ and irreproducibility ^32^. To address the research reproducibility problem, we experimented with requiring two separate papers presenting evidence of a causal miRNA-disease relationship rather than just one. While the total pleiotropy of miRNAs decreased somewhat (on average, pleiotropy per miRNA dropped by 54%), the general trends remained the same. Future work should include addressing these limitations through expanded searches and ontology refinements, utilizing more powerful AI models and exploring additional miRNA information resources.

## Conclusion

We have demonstrated the use of a LLM AI to extract causal information from published abstracts regarding miRNA research with similar accuracy to human experts. This allowed us to quantify, for the first time, miRNA pleiotropy, identifying several intrinsic factors that influence a miRNA’s disease impact. Despite the obvious relevance of pleiotropy to translational miRNA research, such questions have not been asked or answered before. Finally, we have suggested understudied miRNAs that likely cause more diseases than currently known. Code and databases, both miRAIDD and IMAPP are freely available. We anticipate many more AI-driven applications, and with superior accuracy, will contribute to the literature in the years to come.

## Data Availability

The miRAIDD database is available at https://github.com/Wanff/miraidd

## Code Availability

Our code is available at https://github.com/Wanff/miraidd

## Author Information

### Affiliations

Harvard University, Cambridge, MA, 02132, USA

R. Wang

Channing Division of Network Medicine, Brigham and Women’s Hospital and Harvard Medical School, Boston, MA, 02115, USA

J. Hecker and M. McGeachie

### Ethics Declarations Competing Interests

The authors declare that they have no competing interests.

## Bibliography

1. Ardekani, A. M. & Naeini, M. M. The Role of MicroRNAs in Human Diseases. Avicenna J. Med. Biotechnol. 2, 161–179 (2010).

2. Bartel, D. P. MicroRNAs: Genomics, Biogenesis, Mechanism, and Function. Cell 116, 281– 297 (2004).

3. Jenike, A. E. & Halushka, M. K. miR-21: a non specific biomarker of all maladies. Biomark. Res. 9, 18 (2021).

4. Huang, Z., et al. HMDD v3.0: a database for experimentally supported human microRNA– disease associations. Nucleic Acids Res. 47, D1013–D1017 (2019).

5. Cui, C., Zhong, B., Fan, R. & Cui, Q. HMDD v4.0: a database for experimentally supported human microRNA-disease associations. Nucleic Acids Res. gkad717 (2023) doi:10.1093/nar/gkad717.

6. Radford, A., et al. Language Models are Unsupervised Multitask Learners.

7. Wei, J. et al. Emergent Abilities of Large Language Models.

8. Introducing ChatGPT. https://openai.com/blog/chatgpt.

9. Kozomara, A., Birgaoanu, M. & Griffiths-Jones, S. miRBase: from microRNA sequences to function. Nucleic Acids Res. 47, D155–D162 (2019).

10. PubMED Advanced Search Builder. National Library of Medicine (US), National Center for Biotechnology Information.

11. Bello, S. M., et al. Disease Ontology: improving and unifying disease annotations across species. Dis. Model. Mech. 11, dmm032839 (2018).

12. Medical Subject Headings (MeSH) SPARQL API. National Library of Medicine (US), National Center for Biotechnology Information.

13. Nan, Y., et al. MiRNA-451 plays a role as tumor suppressor in human glioma cells. Brain Res. 1359, 14–21 (2010).

14. Zhang, Y., Tang, S., Yang, W. & Du, F. let-7b-5p suppresses the proliferation and migration of pulmonary artery smooth muscle cells via down-regulating IGF1. Clinics 77, 100051 (2022).

15. New and improved embedding model. https://openai.com/blog/new-and-improved-embedding-model.

16. Paszke, A., et al. PyTorch: An Imperative Style, High-Performance Deep Learning Library. Preprint at http://arxiv.org/abs/1912.01703 (2019).

17. Panwar, B., Omenn, G. S. & Guan, Y. miRmine: a database of human miRNA expression profiles. Bioinformatics 33, 1554–1560 (2017).

18. Ru, Y., et al. The multiMiR R package and database: integration of microRNA–target interactions along with their disease and drug associations. Nucleic Acids Res. 42, e133 (2014).

19. Enright, A. J., et al. MicroRNA targets in Drosophila. Genome Biol. 5, R1 (2003).

20. McGeary, S. E., et al. The biochemical basis of microRNA targeting efficacy. Science 366, eaav1741 (2019).

21. Chen, Y. & Wang, X. miRDB: an online database for prediction of functional microRNA targets. Nucleic Acids Res. 48, D127–D131 (2020).

22. Wang, K. R. & McGeachie, M. J. DisiMiR: Predicting Pathogenic miRNAs Using Network Influence and miRNA Conservation. Non-Coding RNA 8, 45 (2022).

23. R Core Team. R: A language and environment for statistical computing. R Foundation for Statistical Computing (2021).

24. McInnes, L., Healy, J. & Melville, J. UMAP: Uniform Manifold Approximation and Projection for Dimension Reduction. Preprint at http://arxiv.org/abs/1802.03426 (2020).

25. Vallat, R. Pingouin: statistics in Python. J. Open Source Softw. 3, 1026 (2018).

26. Gao, Y., Jia, K., Shi, J., Zhou, Y. & Cui, Q. A Computational Model to Predict the Causal miRNAs for Diseases. Front. Genet. 10, (2019).

27. Kim, T. & Croce, C. M. MicroRNA: trends in clinical trials of cancer diagnosis and therapy strategies. Exp. Mol. Med. 55, 1314–1321 (2023).

28. Godard, P. & van Eyll, J. Pathway analysis from lists of microRNAs: common pitfalls and alternative strategy. Nucleic Acids Res. 43, 3490–3497 (2015).

29. Ratwani, R. M., Bates, D. W. & Classen, D. C. Patient Safety and Artificial Intelligence in Clinical Care. JAMA Health Forum 5, e235514 (2024).

30. Karkera, N., Acharya, S. & Palaniappan, S. K. Leveraging pre-trained language models for mining microbiome-disease relationships. BMC Bioinformatics 24, 290 (2023).

31. Easterbrook, P. J., Gopalan, R., Berlin, J. A. & Matthews, D. R. Publication bias in clinical research. The Lancet 337, 867–872 (1991).

32. Resnik, D. B. & Shamoo, A. E. Reproducibility and Research Integrity. Account. Res. 24, 116–123 (2017).

