## Supplemental Methods for "Quantifying the massive pleiotropy of microRNA: a human microRNA-disease causal association database generated with ChatGPT"

Supplemental Materials

**Supplemental Methods**

**ChatGPT Prompt**

*You are an intelligent AI assistant who, given an abstract, determines whether a given miRNA in that abstract plays a causal role in the described disease.*

*Here are some things you should know:*

*Abstracts that mention causal intervention experiments like transfection of miR mimic, knockout/knockdown experiments, and rescue expression experiments tend to be causal.*

*Abstracts that use differential expression analysis, computational methods, or identify biomarkers tend to be not causal.*

*If a miRNA causally influences a protein/gene that is itself causal for the disease, the miRNA is also causal.*

*If a causal protein/gene influences a miRNA, that miRNA is not necessarily causal.*

*Here's an example task completion:*

*Title: MiRNA-451 plays a role as tumor suppressor in human glioma cells*

*Abstract: MicroRNAs (miRNAs) are small non-coding RNAs that negatively regulate gene expression at the post-transcriptional and/or translational level by binding loosely complimentary sequences in the 3'untranslated regions (UTRs) of target mRNAs. Increased expressions of several miRNAs, specifically hsa-miR-21, have been reported to modulate glioma development. Here we report downregulation of miR-451 in A172, LN229 and U251 human glioblastoma cells. Increased expression of miR-451 by administration of miR-451 mimics oligonucleotides reversed the biology of each of the three cell lines, inhibiting cell growth, inducing G0/G1 phase arrest and increasing cell apoptosis. Further, treatment with miR-451 mimics oligonucleotides diminished the invasive capacity of these cells, as the number of cells invading through matrigel was significantly decreased. Akt1, CyclinD1, MMP-2, MMP-9 and Bcl-2 protein expression decreased, and p27 expression increased in a dose-dependent manner with miR-451 mimics oligonucleotides. Taken together, these studies reveal miR-451 impacts glioblastoma cell proliferation, invasion and apoptosis, perhaps via regulation of the PI3K/AKT signaling pathway. We propose an essential role for miR-451 as a tumor-suppressor of human glioma.*

*Question: Does mir-451 play a causal role in the disease described above?*

*Your Answer: Since the authors used mir-451 mimic to decrease the invasive capacity of glioblastoma cells, mir-451 causes glioma.*

*Think step by step and use your best judgement to determine causal miRNAs from abstracts. In your response, be as succinct as possible and mention the disease in your response.*

*Title: [[Title]]*

*Abstract: [[Abstract text]]*

*Question: Does [[miRNA-Name]] play a causal role in the disease described above?*

*Your Answer:*

**List of Cancer MeSH Terms**

MeSH terms containing one of these strings were considered to be cancer-related: "carcinoma”, "neoplasms”, "cancer”, "tumor” "leukemia" "lymphoma", "polycythemia vera", "blastoma", "sarcoma"
